# Glutamic Acid at position 168 is a constitutive activator of Tank Binding Kinase 1 catalytic function

**DOI:** 10.1101/2025.08.17.669475

**Authors:** Noopur Bhore, Anubhuti Sarkar, Zhi Yao, Susanne Herbst, Patrick A. Lewis

**Author notes:** Equal contribution (Noopur Bhore, Anubhuti Sarkar).

## Abstract

TANK binding kinase 1 (TBK1) is serine/threonine protein kinase member of the inhibitor of nuclear factor-κB kinase family, with links to the etiology of familial as well as idiopathic Amyotrophic Lateral Sclerosis. It contributes to several regulatory cellular processes such as autophagy, inflammation and apoptosis. Reduction or loss of TBK1 kinase activity is associated with increased risk of ALS, and so understanding the molecular basis of this activity is an important research priority. In this current study, the role of the E168 residue, located adjacent to the active site of TBK1, has been assessed using a combination of artificial and naturally occurring variants found at this codon – evaluated using multiple readouts for TBK1 kinase activity. The results suggest that the negative charge resulting from the presence of a glutamic acid at this codon is a constitutive activator of TBK1 activity.

## Introduction

Amyotrophic Lateral Sclerosis (ALS) is a progressive, fatal, and multifactorial neurodegenerative disorder characterized by the selective degeneration of upper and lower motor neurons, resulting in the denervation and subsequent atrophy of skeletal muscle fibers, leading to progressive motor impairment and respiratory failure, with a median survival age of 2–5 years following diagnosis (Chio et al. 2009; Sacks et al. 2022; Peters et al. 2015). Mendelian Studies have identified multiple loci with a strong predisposition linked to ALS, including hexanucleotide expansions in chromosome 9 open reading frame 72 (*C9orf72*),and mutations in superoxide dismutase 1 *(SOD1)*, TAR DNA-binding protein 43 *(TARDBP)*, fused in sarcoma *(FUS)*, Optineurin *(OPTN)* and TANK-binding kinase 1 *(TBK1)* (Renton et al. 2011; Rosen et al. 1993; Sreedharan et al. 2008; Vance et al. 2009; Maruyama et al. 2010; Freischmidt et al. 2015).

TBK1 is a 729-amino acid, multifunction serine/threonine kinase belonging to the inhibitor of nuclear factor-κB (IKK) family, playing a pivotal role in regulating autophagosome-mediated degradation of ubiquitinated cargo and orchestrating inflammatory responses through substrate phosphorylation of autophagy adaptors (Freischmidt et al. 2015; Oakes et al. 2017). TBK1 is highly expressed in neuronal and glial cells of brain regions essential for learning and memory, movement, balance and posture, and other executive functions (Oakes et al. 2017). Dysfunction of TBK1 is linked to a number of human diseases, most prominently (ALS) and ALS/frontotemporal dementia (ALS/FTD), but also herpes simplex encephalitis (HSE), diabetes, obesity, cancer, and normal tension glaucoma (NTG) (Ahmad et al. 2016). TBK1 has multiple roles within the cell, activating immunological response to viral or proteostatic stress stimuli, endosomal response, autophagy, or apoptosis (Oakes et al. 2017; Runde et al. 2022; Ahmad et al. 2016; Shao et al. 2022; Talaia et al. 2024; Harding et al. 2021; Fischer et al. 2025; Lu et al. 2021).

Genetic studies have identified over 90 distinct mutations in Tank-Binding Kinase 1 (TBK1) across sporadic and familial ALS cases, highlighting its critical role in disease pathogenesis (Freischmidt et al. 2017; Harding et al. 2021). These mutations span nonsense, frameshift, missense, and single amino acid deletions, each exhibiting distinct functional consequences (Oakes et al. 2017). While nonsense and frameshift mutations significantly disrupt TBK1 expression at both the mRNA and protein levels—suggesting haploinsufficiency as a potential pathogenic mechanism— missense variants and small deletions exert more nuanced effects that may not necessarily reduce protein abundance (Freischmidt et al. 2015; Pottier et al. 2015). ALS-associated TBK1 mutations impair key functional attributes, including dimerization, interaction with mitophagy receptor optineurin (OPTN), autoactivation, and substrate phosphorylation (Li et al. 2016; Ye et al. 2019; de Majo et al. 2018).

Despite extensive investigation, there are a number of aspects of TBK1 enzymatic function, as well as dysfunction in disease, that remain unclear. Using a combination of homology modelling and directed mutagenesis, in this current study the role of a key glutamic acid residue, E168, located in the activation loop of the kinase domain of TBK1, has been investigated to examine its contribution to TBK1 kinase activity.

## Materials and Methods

### Post translational modifications

Data for post translational modification of TBK1, IKKα, IKKβ and IKKε were accessed through the phosphosite portal (Hornbeck et al. 2015).

### Sequence Alignment

The protein sequences of TBK1 (NP_037386.1; 1-729 aa), IKKα (NP_001269.3; 1-745 aa), IKKβ (NP_001547.1; 1-756 aa) and IKKε (NP_054721.1; 1-716 aa) were obtained from the National Center for Biotechnology Information (NCBI) and aligned using the Needleman-Wunsch algorithm (Sayers et al. 2024).

### Structural Modelling

The orientation of E168 of TBK1 was assessed in a crystal structure of a dimer of full-length, active, S172-phosphorylated TBK1(PDB ID: 4IW0) (Larabi et al. 2013). The full-length dimer structure of IKKα phosphorylated at S176 and S180 was modelled using the AlphaFold3 server (Abramson et al. 2024) and the highest confidence structure was used to estimate the orientation of pS176. All structures were displayed using Mol*Viewer (Sehnal et al. 2021).

### Plasmids

The open reading frame of human TBK1 (NM_013254) cloned into pcDNA3.1(+)-C-HA was obtained from GenScript. Point mutations were introduced by site-directed mutagenesis using the Q5®-site directed mutagenesis kit (E0552S, New England Biolabs). The following plasmids were generated – TBK1-E168A, TBK1-E168S, TBK1-E168K, and TBK1-S172A. The plasmid sequences were verified by Sanger sequencing courtesy of Genewiz (Azenta Life Sciences). The plasmids were maintained in *E. coli* DH5α (11583117, Thermo Scientific) and isolated using the QIAprep spin miniprep plasmid extraction kit (27106, QIAgen). For transfections, Fugene® HD reagent (E2311, Promega) was used according to the manufacturer’s instructions in a 1:3 DNA: Reagent ratio. Mock controls were treated with the transfection reagent, but no plasmids were used. Primer sequences used for site-directed mutagenesis are as follows:

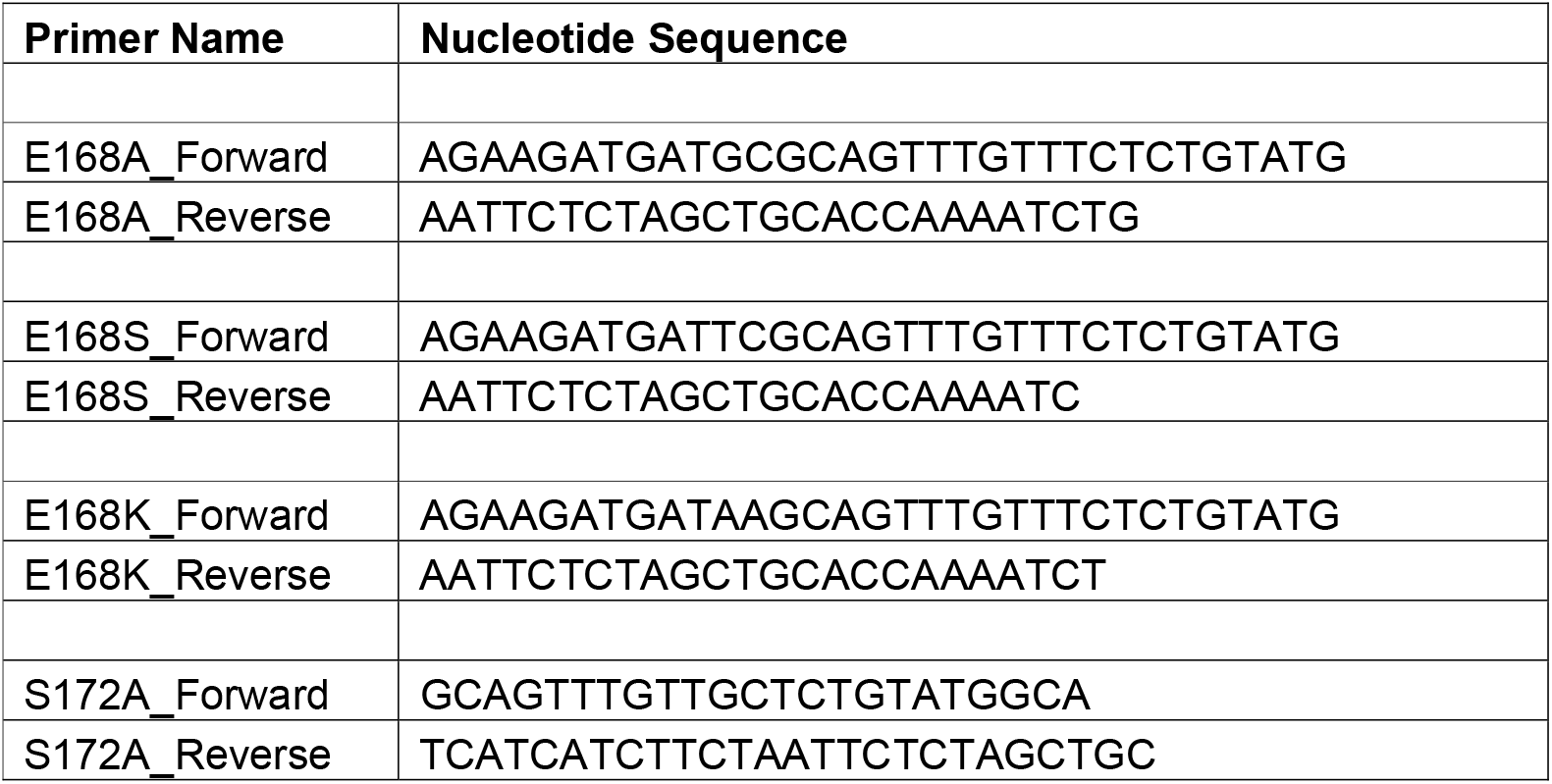

### Cell culture

Human embryonic kidney derived HEK293T/17 (CRL 11268) [hereafter referred as HEK293] adherent cells procured from American Type Culture Collection (ATCC) were cultured in Dulbecco’s Modified Eagle Medium (DMEM) and 10% fetal bovine serum (FBS) (A5670701, Thermo Scientific) without the addition of any antibiotics. Cells were seeded in a 12-well plate for the western blotting experiments at a density of 2.5×10^5^ cells per well. Cells were reverse transfected with the plasmids of interest (1 µg per well) for 48 hours and incubated at 37°C with 5% CO_2_, before being treated with 1µM TBK1-inhibitor GSK8612 (S8872, Sellekchem) in DMEM+10% FBS, for an hour. The cells were then harvested and processed for immunoblotting.

### Immunoblotting

The harvested cells were lysed in a cell lysis buffer (9803S, Cell Signaling Technology) supplemented with Halt™ protease inhibitor cocktail (1861280, Thermo Scientific) and then treated with a sample loading buffer. The loading samples were denatured at 80°C for 8 minutes, centrifuged at 5000x*g* for 1 minute, and the supernatant loaded onto a NuPAGE 4-12% precast gel (10338442, Fisher Scientific) for western blotting. The gels were then transferred onto TransBlot® Turbo™ PVDF membranes (1704156, Bio-Rad) and blocked using 5% milk in 1x TBST (Tris-buffered saline, 0.05% Tween 20) for 1 hour. The blocked membranes were subjected to overnight incubation with the antibody preparation as follows: Anti-phospho TBK1 (D5C2, 5483S, Cell Signaling Technology), Anti-TBK1 (3013S, Cell Signaling Technology), Anti-phospho RAB7 (AB302494, Abcam), Anti-RAB7 (E907E, 95746S, Cell Signaling Technology), Anti-β-actin HRP (A3864, Sigma), and Anti-HA tag (H3663, Sigma). All the primary antibodies were diluted 1:1000. The secondary antibodies used were goat Anti-Rabbit HRP (A0545, Sigma) and goat Anti-Mouse HRP (A3682, Sigma) at a concentration of 1:10,000 in 5% milk prepared with 1x TBST for 1 hour at ambient temperature. The blots were imaged using horse-radish peroxidase chemiluminescent substrate on an iBright™ imager (Thermo Fisher Scientific).

### Statistical Analysis

The immunoblot results were analyzed using GraphPad Prism 10.3.1. All statistics employed an ordinary one-way ANOVA test followed by Dunnett’s multiple comparisons post-hoc test. Graphs represent the mean ± standard error of the mean of 4 independent experiments, where each repeat further comprised of technical duplicates.

## Results

To investigate the phosphoregulation of TBK1, post-translational modification data for each of the IKK family was accessed through the phosphosite portal. This revealed distinct patterns of phosphorylation across the family (**figure S1** and **S2**, and **tables S1-4**). Focusing on phosphorylation of the kinase domains of the IKKs, a divergence between IKKα and IKKβ, and TBK1 and IKKε, was observed – consistent with previous analyses. Dual phosphorylation was observed in IKKα and IKKβ, (at residues S176/S180, and S177/S181 respectively), and a single phosphorylation event in TBK1 and IKKε, at S172 residue – with S176 in IKKα, S177 in IKKβ and S172 in TBK1/IKKε being conserved across the family (**figure 1A**). Intriguingly, both TBK1 and IKKε shared a glutamic acid (E168 in both kinases) at the equivalent residue to the S176/S177 residue in IKKα and IKKβ. Analysis of the crystal structures for TBK1 and a structural model of phosphorylated IKKα revealed that the S176 and E168 residues are spatially homologous (**figure 1B-C**).

**Figure 1.**
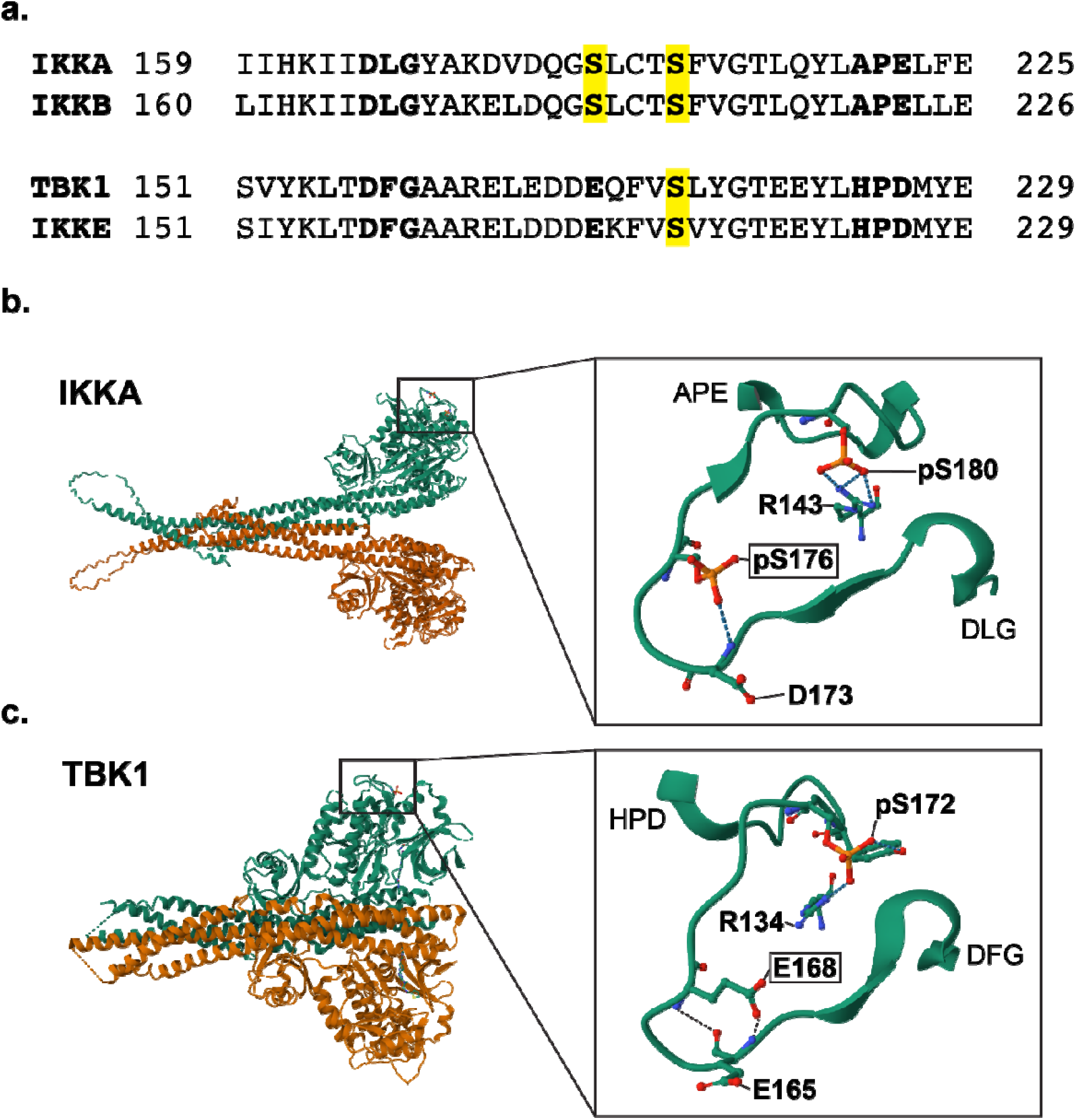
Structural comparison of the activation loop of the IKK family of kinases. (a) Sequence alignment of the activation loop (DLG/DGF motif to APE/HPD motif) of IKKA, IKKB, TBK1 and IKKE. The phosphorylation sites are highlighted in yellow, and the conserved Glutamic acid (E) at position 168 in TBK1 and IKKE is emphasised in bold. (b) AlphaFold3 structural model of S176 and S180 phosphorylated IKKA dimer. The inset shows the activation loop, highlighting the positioning and interactions of the phosphorylated serines. (c) Crystal structure of the S172 phosphorylated TBK1 dimer (PDB ID: 4IW0). The inset shows the activation loop, highlighting the homology in positioning of TBK1 E168 to phosphorylated S176 of IKKA.

Notably, the glutamic acid at this residue in TBK1 provides a negative charge with some functional overlap with phospho-serine in IKKα/β, with the serine residue in the latter providing modifiable phospho-regulation of this location. Previous studies of S176 and S177 in IKKα/β have tested the role of this residue by mutating it to either an alanine (incapable of phosphorylation) or glutamic acid (acting as a phosphomimetic) (Ling et al. 1998; Delhase et al. 1999). These data suggest that in IKKα/β, this residue acts as a key regulator of kinase activity – with a glutamic acid phosphomimetic acting to constitutively activate the kinase activity of these proteins. To examine whether the E168 residue is polymorphic in human populations, variation at this codon was assessed using the Gnomad dataset, revealing a single E168K variant.

To test the biochemical role of the E168 residue, and the consequences of variation at this codon, HA epitope tagged TBK1 constructs for wildtype, E168A (removing the negative charge), E168S (equivalent to the homologous residue in IKKα/β), and E168K were transfected into HEK293 cells. Phosphorylation of TBK1 at S172, as well as Rab7 at S72 as a validated substrate for TBK1, was evaluated by immunoblot (**figure 2**). As expected, inhibition of TBK1 resulted in a dramatic decrease in kinase activity as indicated by phosphorylation of Rab7. Mutation of the S172 residue to an alanine likewise resulted in ablation of kinase activity, as previously reported (Kishore et al. 2002). Examining the impact of the E168 variants, all three significantly reduced TBK1 kinase activity. E168A and E168S lowered phosphorylation of the S172 residue by ≈ 80% compared to wild type, with E168K reducing phosphorylation below the threshold of detection. This was also the case for TBK1 kinase activity directed towards Rab7 S72. These data support a critical role for the E168 residue in regulating the kinase activity of TBK1.

**Figure 2.**
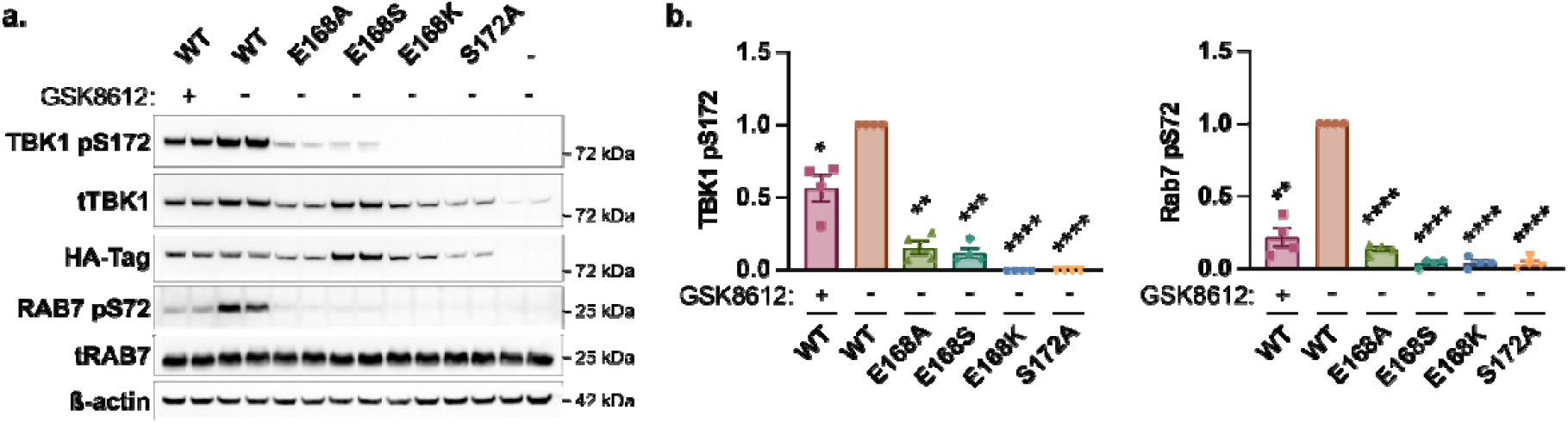
Variation at TBK1 E168 results in TBK1 loss-of-function. HEK293 cells were transfected with the indicated HA-tagged TBK1 constructs, followed by treatment with the TBK1 kinase inhibitor GSK8612 (1µM for 1 hr). (a) TBK1 kinase activity was assessed by Western Blotting for TBK1 pS172 and Rab7 pS72. (b) Graphs represent TBK1 pS172 and Rab7 pS72 levels adjusted for total TBK1 levels and normalised to WT transfected cells. The data shows the mean +/− SEM of 4 independent experiments. Results were compared to WT transfected cells by one-way ANOVA, followed by Dunnett’s multiple comparisons test. ^**^ *p<0*.*01*, ^***^ *p<0*.*001*, ^****^ *p<0*.*0001*.

## Discussion

The *TBK1* gene contributes to risk of developing ALS/FTD as both a monogenic Mendelian locus and *via* common variation as a risk locus. As such, and coupled to its status as a kinase regulator of signal transduction pathways, it is a strong candidate for modulation as a drug target for these disorders. However, the gaps in our understanding of how TBK1 functions with regard to its enzymatic function and the complexities of its signaling network present substantial obstacles to drug development. There is a particular medicinal chemistry challenge with regard to the genetic data supporting a boosting of TBK1 activity as being potentially beneficial, requiring small molecule allosteric activators rather than inhibitors. In order to better understand the activation of TBK1, in this study a comparative analysis of post-translational modification across the IKK family was carried out, identifying a key glutamic acid residue at codon 168 in TBK1 and IKKε that is homologous to a modifiable – and phosphorylated – serine in IKKα and IKKβ. This residue sits within the activation loop of TBK1, critical to kinase regulation and activity (Adams 2003; Reinhardt and Leonard 2023). Based upon this homology, the role of the E168 residue was tested by mutating it to alanine and serine, as well as to a lysine variant identified in the Gnomad dataset. All three variants significantly reduced TBK1 kinase activity as measured by autophosphorylation, suggesting that the negative charge imparted by the side chain of glutamic acid is a required for the kinase function of TBK1. These data also suggest that naturally occurring loss-of-function variants at this residue, such as the E168K variant, may act as a risk factor for ALS/FTD. The results of this study highlight the critical role of residues around the active site of TBK1 for activity, and supports further investigation as to whether targeting this residue could facilitate modulation of TBK1 kinase.

## Acknowledgements

This study was funded by the My Name’5 Doddie Foundation (grant MN5DF/CAAu23/100008). PAL is a Royal Society Industry research fellow in partnership with LifeArc (IF\R2\222002).

## Author Contributions

PAL conceptualized the study. NB, SH and PAL designed the experiments. AS, NB, SH, and PAL performed the experiments, analyzed the data, and wrote the manuscript. All authors edited the manuscript and approved the final draft for publication.

## Abbreviations

A: Alanine amino acid
E: Glutamate amino acid
K: Lysine amino acid
S: Serine amino acid
ALS: Amyotrophic lateral Sclerosis
C9orf72: Chromosome 9 Open Reading Frame 72
FTD: Frontotemporal dementia
FUS: Fused in sarcoma
GoF: Gain of function mutations
HEK293: Human embryonic kidney cells
IKK: Inhibitor of nuclear factor kappa-B
LoF: Loss of function mutations
Nf-κB: Nuclear Factor Kappa-B
p-: phosphorylated
p62: Phosphotyrosine-Independent Ligand For The Lck SH2 Domain Of 62 KDa
RAB7: Ras-related protein
TBK1: TANK-binding kinase 1
TDP-43: TAR DNA-binding protein 43

## Notes

### Competing Interest Statement

The authors have declared no competing interest.

